# Flat-field illumination for quantitative fluorescence imaging

**DOI:** 10.1101/297242

**Authors:** Ian Khaw, Benjamin Croop, Jialei Tang, Anna Möhl, Ulrike Fuchs, Kyu Young Han

**Affiliations:** CREOL, The College of Optics and Photonics, University of Central Florida, Orlando, Florida, USA.; asphericon GmbH, Stockholmer Str. 9, D-07745 Jena, Germany

## Abstract

The uneven illumination of a Gaussian profile makes quantitative analysis highly challenging in laser-based wide-field fluorescence microscopy. Here we present flat-field illumination (FFI) where the Gaussian beam is reshaped into a uniform flat-top profile using a high-precision refractive optical component. The long working distance and high spatial coherence of FFI allows us to accomplish uniform epi and TIRF illumination for multi-color single-molecule imaging. In addition, high-throughput borderless imaging is demonstrated with minimal image overlap.

## Introduction

Wide-field fluorescence imaging is a powerful tool for studying molecular mechanisms of subcellular processes. It is widely used in screening drugs for various diseases, and in profiling cellular phenotypes by an image-based high-throughput system^1,2^. In these applications, the fluorescence intensity is the primary information for the analysis of images. Since the emitted fluorescent signal from a fluorophore is dependent on the amount of incident light, a uniform illumination is critical to perform quantitative analysis. Many factors including light sources and illumination optics contribute the uniformity of the illumination. For instance, in epifluorescence excitation, a lamp or light-emitting diode is used as a light source where a traditional Köhler illuminator or microlens array^3^ creates the uniform illumination.

For imaging single-molecules^4^ or organelles^5^ near the cell surface, a laser is utilized as an excitation source due to its high brightness and high spatial coherence, however the Gaussian illumination profile of the laser has several drawbacks. The uneven illumination of a Gaussian-shaped beam distorts the intensity data and makes detailed analysis challenging. Moreover photobleaching makes it more complicated to analyze the image because it results in a permanent loss of fluorescence signal and the bleaching time strongly depends on the illumination intensity^6^. When revealing the stoichiometry of protein complexes using single-molecule fluorescence imaging, the intensity analysis is sensitive to an uneven intensity distribution and therefore counting of the photobleaching steps is mainly used^7,8^, whereas in freely diffusing single-molecule experiments intensity-based analysis is commonly employed to examine the oligomerization of the molecular complex^9,10^. Importantly, in fluorescence nanoscopy the uneven illumination leads to position-dependent resolution and limits the field-of-view (FOV), which is problematic in high-quality super-resolution imaging^11,12^. For example, in single-molecule-based localization imaging^13–15^ the localization accuracy is determined by the number of the emitted photons, which depends on the excitation intensity. In parallelized stimulated emission depletion (STED) microscopy^11^, which exhibits a higher imaging speed compared to conventional single-spot scanning STED^16^, the spatial resolution depends on the inverse square root of the illumination intensity^17^, and as such there is a wide variance in spatial resolution, i.e. 30 nm at the center but ~100 nm at the edge of an imaging area.

Another instance where uneven illumination can be problematic is when performing high-throughput and high-content imaging^18^, where a large FOV on the scale of several hundreds of microns or millimeters is desired. To achieve such a FOV while maintaining high resolution, a grid of images is acquired such that the borders of one image overlap with the adjacent image area, and then the images are stitched together in post-processing. If the illumination is not uniform, the final stitched image will have dimmed borders around each individual image unless a large image overlap is used. A large overlap will either increase the acquisition time or decrease the overall FOV, depending on the number of images taken. Additionally, nonuniform illumination can give unreliable measurements in cell and tissue samples. A flat-field correction is possible to amend the biased intensity through the post-processing of images^18^ but is sensitive to changes to the experimental setup and uncontrollable intensity fluctuations, and is not feasible in low-light applications^19^ such as single-molecule imaging.

A number of approaches have been demonstrated to create a uniform illumination profile in laser-based wide-field fluorescence microscopy. For example, using the centermost portion of the expanded Gaussian beam is the most common approach^20^ but is not ideal because this results in a severe loss of the incident laser power. Other approaches include using a pair of microlens arrays^12,21^ or a multimode fiber combined with a speckle reducer^22,23^, but they still produce inhomogeneous illumination and/or are not suitable for total internal reflection fluorescence (TIRF) microscopy^5^ because they degrade the spatial coherence of the beam making it impossible to focus tightly to the back focal plane of the objective for TIRF illumination.

An attractive method of generating uniform illumination is by utilizing a pair of aspheric lenses^24^ where the first lens redistributes the Gaussian beam uniformly and the second lens re-collimates it, yielding a flat-field illumination (FFI) profile (Fig. 1a). However, this refractive laser beam shaping system has been challenging to employ in fluorescence microscopy due to the requirement of high surface quality and the limited working distance^25^. Here, we overcame these problems using a specially designed beam shaper optimized for a regular fluorescence microscope over a range of wavelengths. In this work we characterize the beam-shaping performance and compare it with optical simulations. We demonstrate multicolor imaging with epi-illumination, quantitative intensity-based single-molecule imaging with TIRF illumination, as well as borderless high-throughput imaging with minimal overlap of stitched images. Our simple and efficient flat-field illumination is a promising approach for quantitative fluorescence imaging.

**Figure 1.**
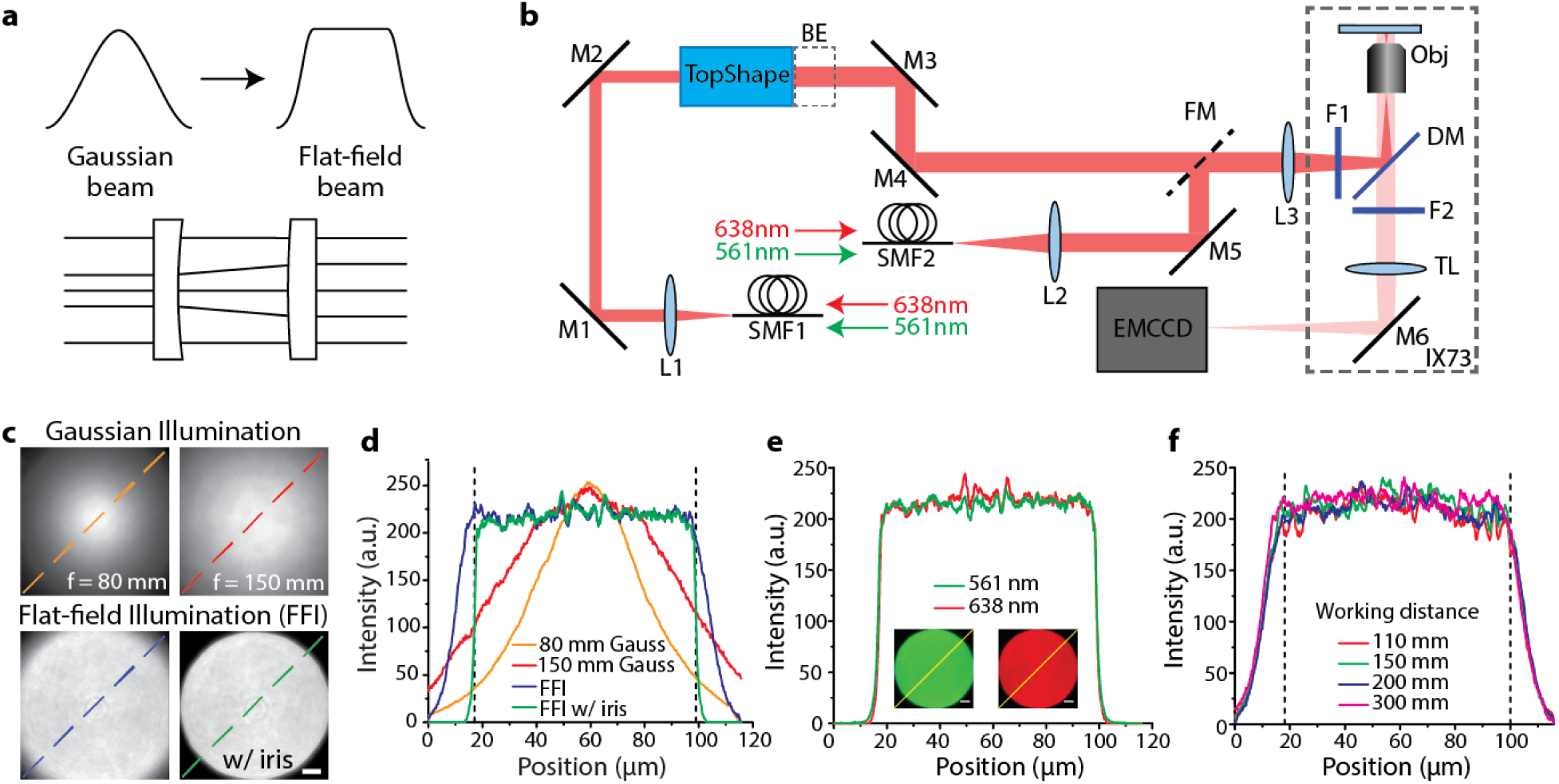
Experimental characterization of FFI. (a) Beam reshaping schematic. (b) Experimental setup. BE, 1.5x beam expander; DM, dichroic mirror; F1-2, excitation/emission filters; FM, flip mirror; L1-3, lenses; M1-6, mirrors; Obj, objective; SMF1-2, single mode fibers; TL, tube lens. (c) Beam profiles of Gaussian beams collimated by a lens with 80 mm or 150 mm focal length and FFI beams without and with an iris. (d) Lineouts taken from the beam profiles in (c) along dashed lines. Vertical dashed lines in (d) indicate a detection region of a camera. (e) Excitation wavelength dependence of FFI. Lineouts taken from multicolor images (inserts) with an iris. (f) Working distance dependence of FFI with 638 nm laser. Scale bars, 10 µm.

## Results

### Flat-field illumination

We first recorded intensity profiles by exciting a thin dye layer with epi-illumination. The laser beam (λ = 638 nm) exiting from a single mode fiber was collimated by an achromatic lens, and FFI was generated by sending the collimated Gaussian beam with a diameter of ~10 mm to a beam shaper called TopShape (Fig. 1b, see Methods section). As shown in Fig. 1c and 1d, FFI generated a flat-top profile, and the full-width at 90% of maximum (FW90M) of FFI was 81.5 µm, which is similar to the diameter of our FOV (~82 µm). For reference, we measured intensity profiles using Gaussian beams collimated by a lens with 80 mm or 150 mm focal length, and their FW90M was 15.0 µm or 28.6 µm, respectively. We estimated the irregularity of the FFI by calculating the root-mean-square of the intensity and it exhibited 2.9% variation. This level of non-uniformity is known not to affect single-molecule imaging and fluorescence nanoscopy^12^. The illumination efficiency (ƞ) of Gaussian beams was 92.9% and 51.4% for 80 mm and 150 mm focal length lenses whereas that of FFI was 85%, implying that the majority of FFI was collected by the detector (see ‘Beam profile measurement’ in Methods section). Multicolor imaging at excitation wavelengths of 561 nm and 638 nm was readily attainable without an additional fiber or collimator (Fig. 1e). The uniform beam profile was maintained over a working distance up to 300 mm at both wavelengths (Fig. 1f, Supplementary Fig. 1). The temporal coherence length of light sources did not affect the illumination profile based on our measurement with HeNe and diode lasers (data not shown here).

### Simulated beam intensity distribution

Since the beam-shaping element (TopShape) is based on refraction, which causes a smooth redistribution of the beam intensity, the flat-top beam profile simulations can be performed using methods based on geometrical optics^26^. Here, the beam profile was calculated at a certain distance behind the beam shaping system before entering the microscope. In Fig. 2a the normalized intensity distribution at a wavelength of 640 nm is shown for different working distances. In general, a constant flat-top beam profile was generated for distances up to 600 mm. However, at a working distance of 600 mm the intensity distribution has slight inhomogeneity because the beam-shaping system used was optimized for shorter working distances. Beam shaping is based on manipulating the phase front relation of the incoming laser beam to achieve the desired output distribution within a certain range. Beyond this region the beam profiles show higher variations due to the increasing mismatch in phase relation. The length of this region depends on the design of the beam shaping device. In the case of the current beam shaping system, the addressed working distance is limited to about ~300 mm for a plateau homogeneity of more than 95%. Note that our experimental beam profiles in Fig. 1d and 1e are measured at a working distance of 300 mm. The working distance range for a very homogeneous flat-top beam profile can be extended by using an additional beam expander behind the beam shaping system, as shown in Fig. 2b. Another simulation was carried out concerning the wavelength dependence of the beam shaping system at a fixed working distance (300 mm). As shown in Fig. 2c, there was nearly no difference in performance between three different laser wavelengths (488 nm, 561 nm and 640 nm).

**Figure 2.**
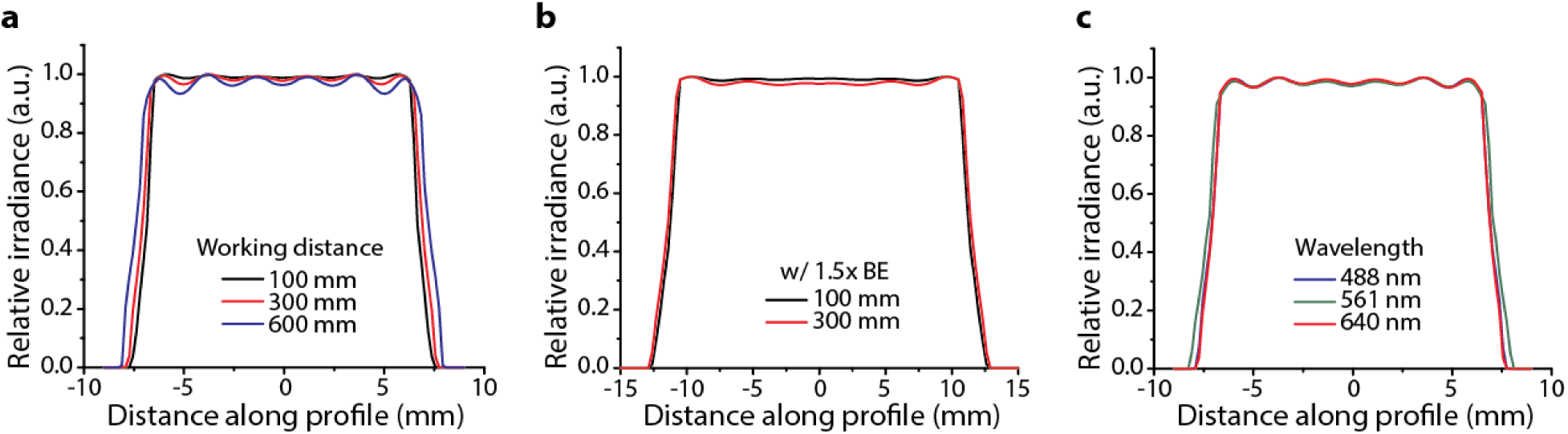
Simulated flat-top intensity distributions. Working distance dependence at a system wavelength of 640 nm without (a) and with (b) an additional beam expander (BE). (c) Excitation wavelength dependence at a working distance of 300 mm.

### Single-molecule imaging and intensity-based analysis

Next, we performed single-molecule imaging on DNA labeled with AlexaFluor 647 (AF647) on a coverslip with objective TIRF microscopy. When subjected to Gaussian illumination, the molecules closest to the center of the beam fluoresced more strongly than those near the periphery, while under FFI the molecules exhibited much more uniform fluorescence (Fig. 3a). The intensity trace of single-molecules at different x-coordinates clearly showed this feature in Fig. 3b. This uniform fluorescence signal yields desirable features in the single-molecule analysis. Firstly, it makes the spot-finding less sensitive to the thresholding value. The thresholding parameter is used to determine if a spot is bright enough to be discerned from the noise level, and generally a global threshold is used^8^. As the normalized threshold value increased, the number of single-molecules was counted and plotted for both Gaussian illumination and FFI. As shown in Fig. 3c, under Gaussian illumination the number of detected spots steadily decreases with an increase to the threshold parameter, while FFI exhibited a noticeable plateau in which the number of spots is insensitive to the thresholding parameter and the true number of spots can be determined.

**Figure 3.**
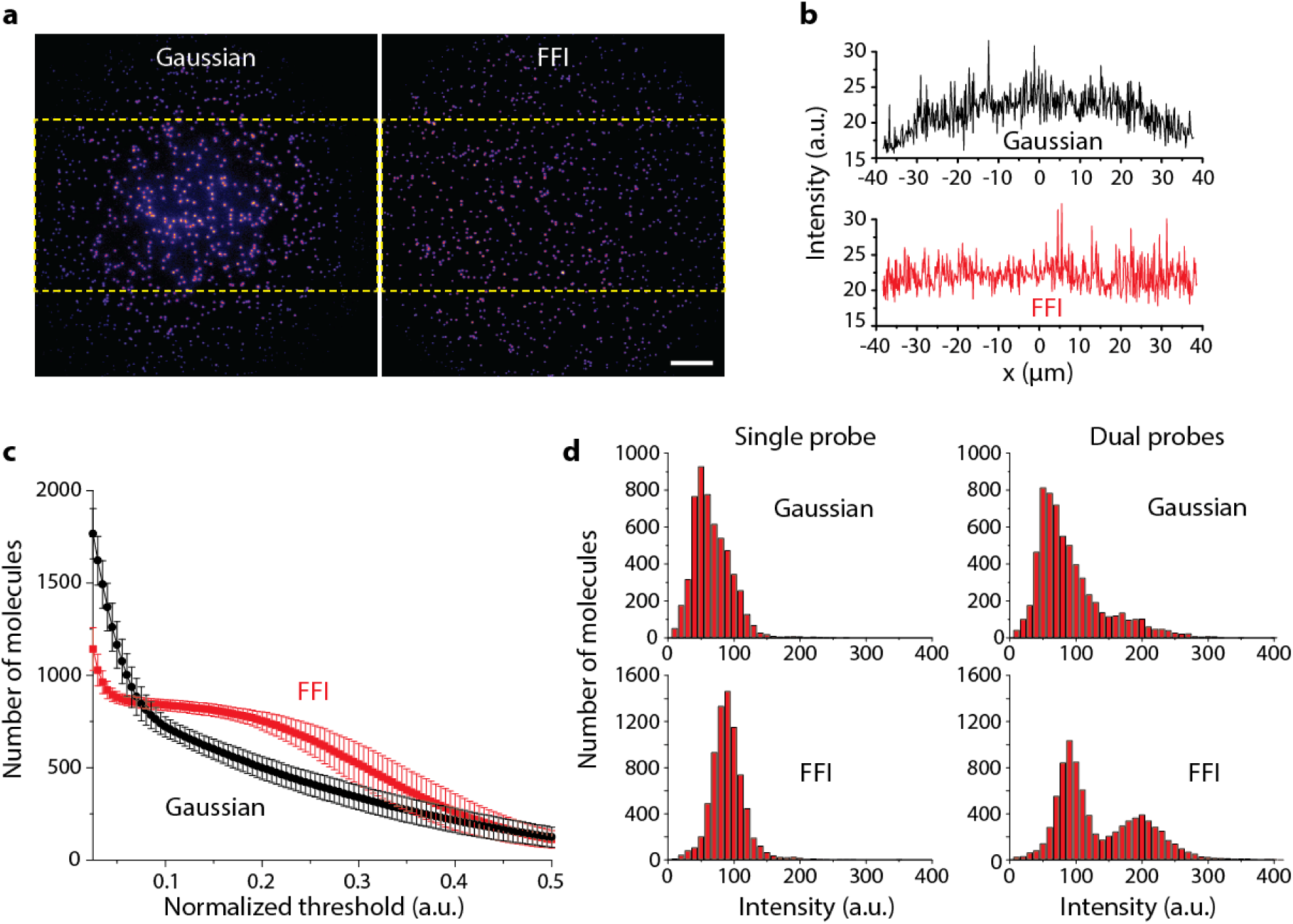
Quantitative single-molecule imaging analysis with FFI. (a) Representative single-molecule images taken under Gaussian illumination and FFI. Scale bar, 10 µm. (b) 1D intensity data taken from the boxed region of (a). (c) Threshold curve showing the dependence of the average number of detected molecules on the background thresholding parameter. Error bars represent the standard deviation from the mean. (d) Intensity distributions of single and dual probe samples imaged under Gaussian illumination and FFI. Images taken from 20 different regions were used for each analysis.

Secondly, we tested whether FFI can improve quantitative intensity-based analysis. For this, we prepared single-probe and dual-probe samples where a capture oligo was hybridized with either one or two complementary probes labeled with AF647. In the dual probe sample, the hybridization process yielded a mixture where the capture oligo bound one probe or two probes. The single probe sample yielded a broadened and skewed distribution under Gaussian illumination (Fig. 3d). By contrast, the intensity distribution by FFI was much narrower and had an identifiable central intensity from which the distribution was well fit to a Gaussian curve. For comparative metrics we calculated the coefficient of variation, i.e. the standard deviation divided by mean, and it was 47% and 31% for the Gaussian illumination and FFI, respectively, meaning that the latter has much less variability in fluorescence intensity. The applicability of FFI for quantitative intensity-based analysis was clearly demonstrated by the dual-probe sample. Figure 3d shows a second population at about twice the intensity of the single probe distribution, which is obscured when subjected to Gaussian illumination. By virtue of the Gaussian distribution for FFI, it was possible to extract the relative population of each species and they were 73% and 27% for one and two probes, respectively. It was straightforward to obtain multi-color single-molecule imaging (Supplementary Fig. 2).

Thirdly, we recorded the photobleaching time under Gaussian illumination and FFI with an elevated excitation power (Fig. 4). The photobleaching time was found to be extremely uniform when the sample was subjected to FFI, whereas under Gaussian illumination single-molecules photobleached more quickly in the center than near the periphery.

**Figure 4.**
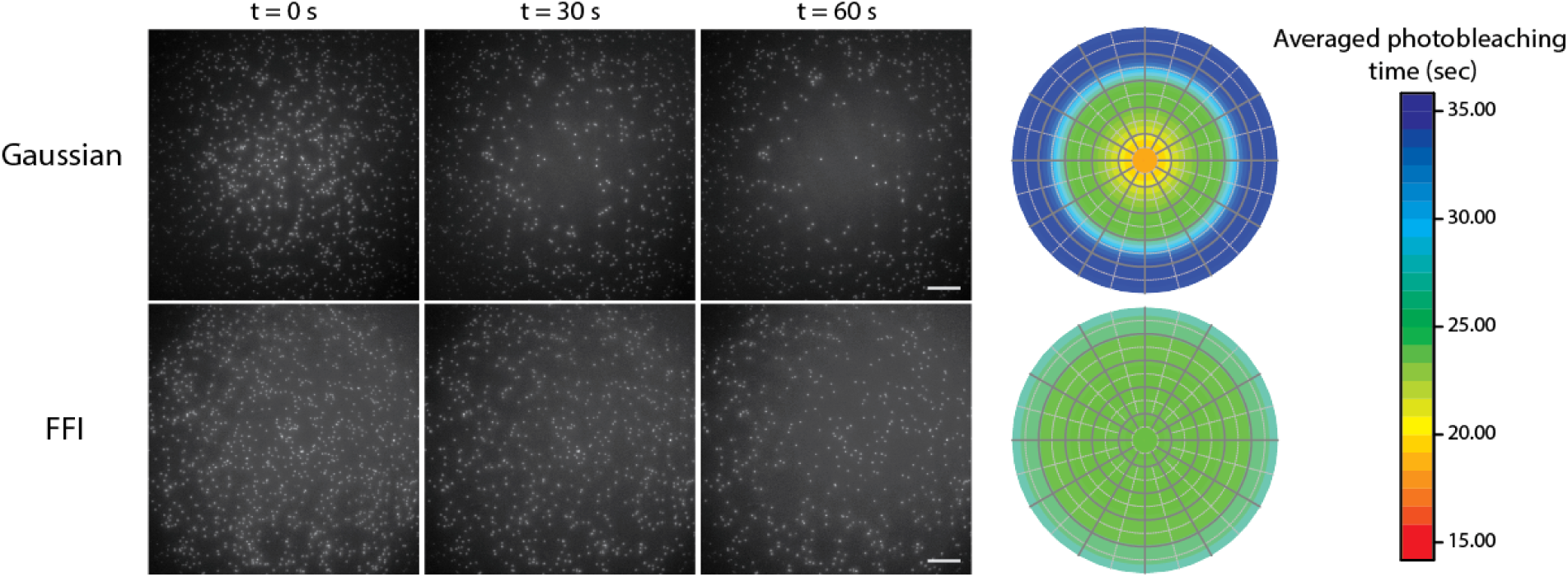
Uniform photobleaching via FFI. Representative single-molecule images taken at times of 0, 30, and 60 seconds under Gaussian illumination and FFI. Colormap showing the average photobleaching time analyzed with images taken from 10 different regions. Scale bars, 10 µm.

### Background suppression in TIRF illumination

We next examined whether FFI effectively eliminated background fluorescence via TIRF illumination by imaging single-molecules in the presence of 5 nM fluorescently labeled molecules. For comparison, we performed the experiment with a multi-mode fiber (MMF) combined with a speckle scrambler that generated a homogeneous profile with epi-illumination. While the FFI provided by the TopShape was able to fully suppress the background, as evident in Fig. 5, the MMF approach showed a highly elevated background level indicating that only partial TIRF was achieved. The measured signal to background ratio was 12.5 ± 7.8 and 5.1 ± 4.1 (mean ± S.D.; *n* = 50) for FFI and MMF, respectively. In our TIRF microscope, an incident beam needed to be focused to a diameter of <120 μm at the back focal plane of the objective for all incident light to contribute to the generation of an evanescent field, however the limited spatial coherence of the MMF prevented tight focusing.

**Figure 5.**
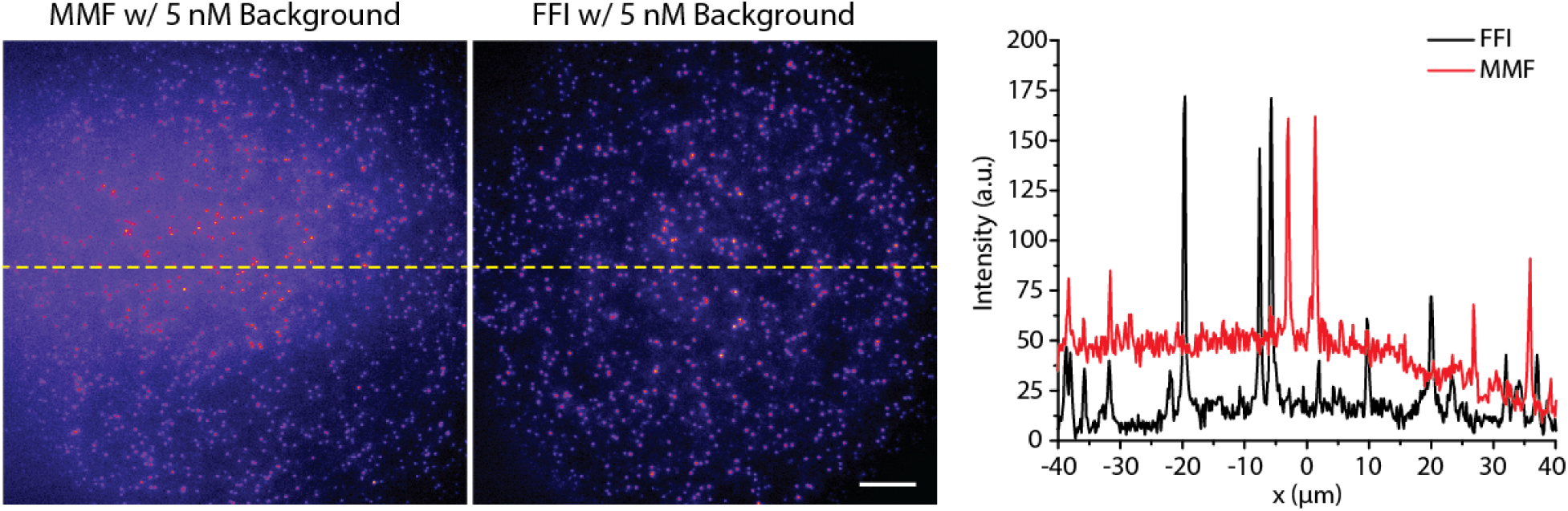
Background reduction by TIRF illumination. Representative images taken under illumination by a multimode fiber (MMF) combined with a speckle scrambler and under FFI in the presence of 5 nM background. Lineouts taken as indicated by the dashed yellow lines show the MMF only achieved partial TIRF as evident by the elevated background level, while FFI achieves full TIRF as seen by the minimal background. Scale bar, 10 µm.

### High-throughput imaging

To examine the potential application of FFI for high-throughput imaging, we acquired a grid of images from mammalian cells with epi-illumination using excitation wavelengths of 561 nm and 638 nm. We used a 1.5x beam expander just after the TopShape (Fig. 1b) to fully cover our camera and the illumination profile was not affected by this additional optical component (Supplementary Fig. 3). As depicted in Fig. 6a, FFI enabled borderless stitched imaging with minimal image overlap (5%), whereas the stitched image using Gaussian illumination had distinct dark borders between imaging areas. This shows that our illumination scheme will increase the imaging speed and minimize photobleaching for high-throughput imaging. In addition, we demonstrated the feasibility of using a low magnification (20x) objective in conjunction with FFI (Fig. 6b).

**Figure 6.**
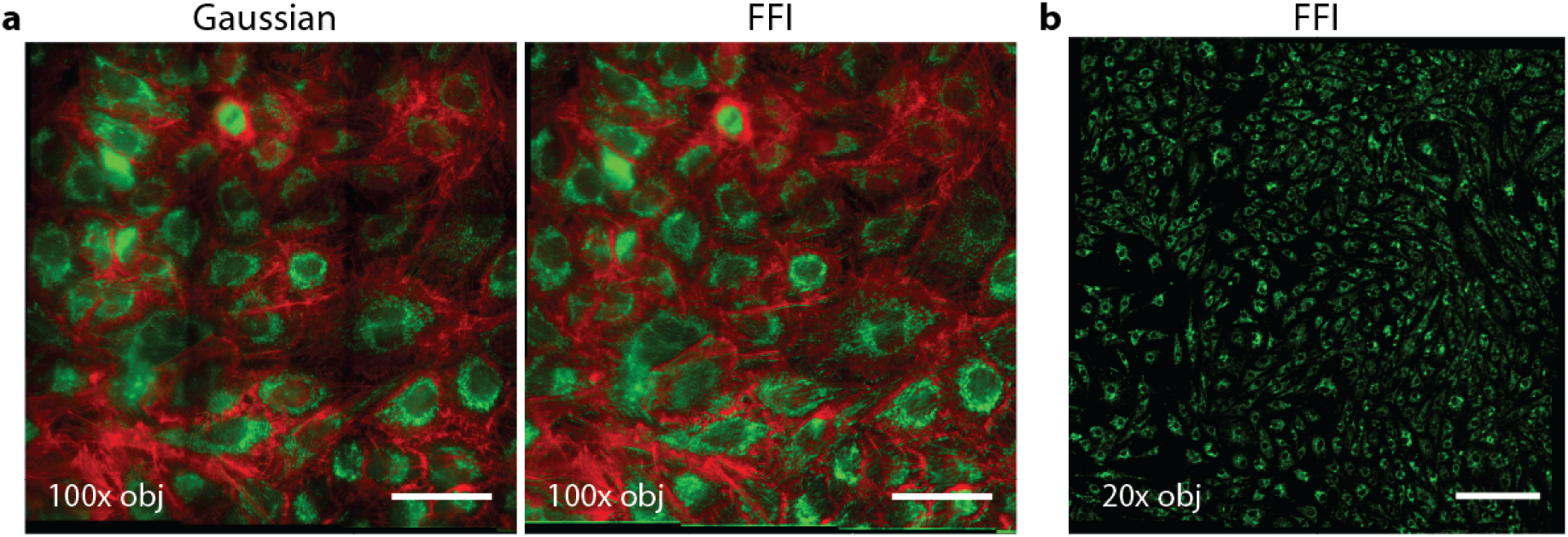
Borderless image stitching under FFI. (a) High-throughput 3×3 multicolor imaging of mitochondria (green) and actin (red) under Gaussian illumination with 150 mm focal length lens and FFI with a 1.5x beam expander with 5% overlap. (b) 3×3 stitched image of mitochondria under FFI with 1.5x beam expander and a 20x objective (10% overlap). Scale bars, 50 µm (a) and 200 µm (b).

## Discussion

We demonstrated flat-field illumination for multi-color wide-field fluorescence microscopy using a refraction-based beam shaping system. As contrasted with other approaches^12,23^, our method is applicable to TIRF illumination, which effectively rejects background fluorescence. Our beam shaping device is extremely tolerant for variations of the incoming laser beam by accepting ±10% variation, while being achromatic as well. This behavior originates from the well-balanced mapping of the incoming rays to the intended flat-top beam profile in combination with a sophisticated material choice, which decreases the sensitivity to input beam diameter variations. The homogenous illumination profile of FFI will enable not only quantitative single-molecule analysis based on intensity information^20^, but also high quality super-resolution imaging with a uniform spatial resolution over a large FOV^11,12^. Further, FFI can potentially be used with a large area detector such as scientific complementary metal-oxide semiconductor (sCMOS) camera to increase the throughput^27^. A new optical design generating a square-shaped beam instead of a round one may be feasible, which would yield greater illumination efficiency. Our compact and high precision beam shaper with a large output diameter can be easily implemented to any commercial wide-field microscope, and thus we expect that simple flat-field illumination approach will greatly advance quantitative fluorescence imaging.

## Methods

### Flat-field illumination fluorescence microscope

Our imaging system was constructed with an Olympus IX73 inverted microscope. Two laser sources (06-MLD 638 nm and 06-DPL 561 nm, Cobolt) were combined, split into two fiber couplers and delivered to the microscope. One output beam from a single-mode fiber (P5-630PM-FC-2, Thorlabs) was collimated with an achromatic lens (L1, f = 63.5 mm, #49-780, Edmund Optics) and sent to the beam shaper (TopShape, asphericon GmbH) where the input size was ~10 mm (1/e^2^), but it is generally acceptable in a range between 9.2 mm and 10.8 mm^26^. Two mirrors (M1 and M2) served as four extra degrees of freedom to guide the beam into the TopShape. The flat-top beam was redirected by mirrors towards the microscope and was focused by a TIR lens (L3, f = 300 mm, AC508-300-A, Thorlabs) to the back focal plane of the objective (UPlanSApo, 100x/1.40 oil, Olympus). To measure Gaussian beam profiles, the other output beam from a single-mode fiber (P5-630A-PCAPC-1, Thorlabs) was collimated with a lens (L2, f = 80 mm or 150 mm, Thorlabs) and sent to the microscope via a flipping mirror which was installed between the TopShape and TIR lens. The TIR lens was mounted on a xyz translator to adjust the incidence angle for epi or TIRF illumination. Fluorescence through a filter cube (laser quad-band TRF89901v2, Chroma) was imaged onto an electron-multiplying charge-coupled device (EMCCD) camera (iXon Ultra 897, Andor). The flat-top profile can be achieved even if the beam through the objective lens is either Airy disc-like or coma-like so long as the intensity profile is flat. Inspection of the intensity profile was done via a live feed by the camera using µManager^28^ while adjusting the angle of incidence to the TopShape.

### Beam profile measurement

A thin dye layer was created by sandwiching a microscope slide with a square coverslip (22 mm in length). 2 µL of a ~1 µM dye solution (STAR635 or Cy3B from Abberior or GE Healthcare, respectively) was dropped on the slide, and the square coverslip was pressed so the dye covered the full area of the coverslip. Epoxy was used to seal the edges. Intensity profiles were measured by exciting the layer of dye with a 638 nm or 561 nm laser with the Gaussian or FFI beam at a working distance of up to 300 mm. The recorded profiles were normalized. We used ImageJ^29^ to measure the intensity profile along the diagonal as indicated in Fig. 1c. The irregularity of the FFI was calculated by the standard deviation divided by the mean of the intensity within the FW90M of the profile. To measure the illumination efficiency (*η*) of the FFI, we inserted a 0.5x demagnifier before the EMCCD camera, which allowed us to capture all of the illumination. The intensity of FFI detected by the camera without the 0.5x demagnifier was divided by the total intensity of FFI to calculate the efficiency. The efficiency for Gaussian beams was calculated using similar principles. The difference in calculating the efficiencies of FFI and the Gaussian beam was that the latter illumination was measured using simulated Gaussian beams with parameters taken from experimental data. The illumination efficiencies for Gaussian beams (f = 80 and 150 mm) and FFI were 93%, 51% and 85%, respectively.

### Optical simulation

The experimental set-up was modelled in the optical design software CodeV (Synopsys). The source is fiber-coupled, whereby the divergence of the laser beam coming out of the fiber depends on the mode field diameter (MFD) of the fiber. This MFD decreases with the wavelength of laser light, which is incorporated in the simulations. The MFD and the divergence angle were calculated for every wavelength, and the fiber output was simulated as a point source, which was collimated by a lens. The distance between the fiber output and the collimation lens was determined by the divergence of the beam and was adjusted aiming for a collimated beam with a diameter of approximately 10 mm at 1/e^2^, which is the required input beam size for our beam shaping system. The collimated beam passed the beam shaping system and the resulting flat-top beam profile was calculated in different distances behind the beam shaping system, which was the working distance. Depending on the simulations performed, an additional beam expander was inserted, to enlarge the output beam by a factor of 1.5. The maximum possible working distance over which the flat-top profile is maintained increases by almost a factor of two. The output intensity distribution was detected using the illumination analysis tool (LUM) in CodeV. The illumination analysis was used to compute the illuminance (or irradiance) distribution across a receiver surface resulting from the luminance of specified 2D or 3D sources. Monte Carlo ray tracing was used to model the transfer of radiation from extended sources to a receiver surface, which was the image surface in this case.

### Preparation of dye-labeled DNA samples

For single-molecule experiments in Fig. 3a-c, Fig. 4, and Fig. 5, biotinylated single-stranded DNA (ssDNA) called oligo 1 was dissolved in MilliQ water for a final concentration of 1 mM. 200 nmoles of AlexaFluor 647 NHS ester (Thermo Fisher) dissolved in anhydrous dimethyl sulfoxide, and 10 nmoles of oligo 1 were mixed in labeling buffer (0.1 M sodium tetraborate pH 8.5). After incubating overnight at room temperature, the sample was purified by ethanol precipitation. The labeled ssDNA was dissolved in 100 µL of T50 (10 mM Tris pH 8 and 50 mM NaCl) buffer, the concentration and labeling efficiency were measured using NanoDrop, and stored at -20°C until use. For dual-probe experiments in Fig. 3d, probes (1 and 2) and capture oligos were diluted in hybridization buffer (200 mM NaCl and 10 mM Tris, pH 8) to a final concentration of 2 µM of each probe and 1 µM of capture oligo, heated at 95 °C for two minutes, then slowly cooled down to room temperature. For single probe experiments, the same procedure was followed but only probe 1 was hybridized with the capture oligo. All chemicals and oligonucleotides were purchased from Sigma-Aldrich and IDT unless specified. The sequence information is listed in Supplementary Table 1.

### Single-molecule fluorescence imaging and analysis

We used biotin-labeled BSA or polyethylene glycol (PEG, Laysan Bio) coated flow chambers as described previously^30^. After washing with T50 buffer, 20 µg/mL neutravidin diluted in T50 was added and incubated for 5 minutes before washing out with T50. ~10 pM of biotinylated DNA labeled with AF647 was incubated for 5 minutes in the flow chamber and washed out. Before imaging acquisition, we added an imaging buffer composed of 2 mM Trolox (Santa Cruz) and an oxygen scavenger (20 mM Tris pH 8.0, 250 mM NaCl, 1 % w/v dextrose, 1 mg/mL glucose oxidase, 0.04 mg/mL catalase). Illumination powers of 4 mW were used for all single-molecule experiments for both Gaussian illumination and FFI, except in the case of the photobleaching experiment. We obtained images from 20 different areas for single-molecule analysis.

A custom MATLAB script was used to identify the location and intensity of each fluorescent spot^30^. For the generation of 1D intensity profile in Fig. 3b, the locations and intensities of molecules in the middle 50% of the field-of-view (as evident by the yellow boxes in Fig. 3a) were stored for both Gaussian illumination and FFI. The intensity of each molecule was plotted as a function of the x-coordinate of the molecule (160 nm precision). In the case where multiple molecules were localized to the same x-coordinate, the intensity of all the molecules at that same coordinate were averaged and plotted as a single point in the 1D profile.

For the generation of the thresholding curve in Fig. 3c, an additional MATLAB script was used in conjunction with the code used for the above intensity distributions. The thresholding parameter was normalized to the intensity of the brightest single molecule in the field-of-view, such that the thresholding parameter was some fractional value multiplied by the intensity of the brightest spot. The x-axis of the threshold curve is the fractional value used to normalize the threshold parameter. The error bars represent the standard deviation from the average number of molecules at each thresholding value. When the threshold is set too low, the spot-finding code incorrectly counts noise as spots, and when it is set too high the code does not count the true single molecule spots and only counts extremely bright spots which are typically impurities or multiple emitters. As the thresholding value is increased past a critical value, the number of spots detected under FFI rapidly drops and the standard deviation increases.

### Photobleaching analysis

DNA labeled with AF647 (oligo 1) was imaged under laser powers of 9 mW for both Gaussian illumination and FFI. For 10 different imaging areas, 2-minute movies were recorded, the locations of molecules detected in the first 10 frames of the movies were stored, and the photobleaching time for all molecules in the movie was measured. The data across all movies was merged, and the illuminated area was divided into six concentric rings, each with a width of 50 pixels, and the average photobleaching time of each ring was calculated. The resulting average photobleaching time was plotted with a colormap.

### Background reduction by flat-field TIRF illumination

We imaged oligo 1 in the presence of 5 nM antibody labeled with AF647 in PEG-passivated flow chambers to check if TIRF illumination was attainable. For the experiment with a multi-mode fiber, we used a bare fiber (Ф = 105 µm, NA 0.15, Draka Prysmian fiber) combined with a vibrating motor (JRF370-18260, ASLONG). We calculated signal-to-background ratio that was defined by (*I*_*S*_ − *I*_*B*_)/*I*_*B*_, where *I*_*S*_ and *I*_*B*_ are the mean of signal and background intensity. 50 single-molecules near the center of the FOV were analyzed.

### Preparation of cell sample

A549 cells (ATCC) were grown on coverslips in a Petri dish with F-12K medium including 10% FBS (F2442, Sigma) and 1% Penicillin/Streptomycin (15140122, Thermo Fisher), incubated in 5% CO_2_ at 37°C for 48 to 72 hours. To stain mitochondria, the cell medium was removed and replenished with pre-warmed staining solution including 50-100 nM MitoTracker Red CMXRos (M7512, Thermo Fisher), incubated in CO_2_ incubator for 30 min. After washing three times with the cell medium, the cells were fixed with 3.7% paraformaldehyde (15710, Electron Microscope Sciences) at room temperature for 15 min. The cells were washed with 1x PBS three times and permeabilized with 0.1% (v/v) Triton X-100 for 5 min. After washing with PBS, the cells were pre-incubated with PBS containing 1% BSA for 30 min, and actins were stained with AF647 phalloidin (A22287, Thermo Fisher) for 20 min. The sample was mounted in Prolong Diamond antifade mountant (P36961, Thermo Fisher) and sealed with epoxy.

### High-throughput Imaging

Using an automated 2D stage (SCAN IM 120 x 80, Marzhauser) controlled by µManager, a 3 × 3 grid was recorded with 638 nm and 561 nm excitation lasers. FFI was expanded 1.5x by installing a beam expander (asphericon GmbH) to the end of TopShape to provide a full FOV illumination to the sample. For comparison, we imaged the same area with a Gaussian beam (f = 150 mm). The red and green images were stitched separately by using the stitching plugin of Fiji software^31^. Parameters for stitching were left as default; 0.30 regression threshold and linear blending for stitching. A leakage of mitochondria signal to the actin image was post-processed. The images collected by FFI did not undergo any prospective or retrospective correction to fix uneven illumination. While stitched images offered seamless transitioning, overlapping of red and green images was not exact due to shifts in image when recording automatically with XY stage.

### Data availability

The data that support the findings of this study are available from the corresponding author upon request.

## Acknowledgments

We thank Rodrigo Amezcua Correa for providing a multi-mode fiber and Jeffrey Moffitt for fruitful discussion regarding a flat-field illumination. This work was supported by DARPA (HR00111720066) and CREOL at University of Central Florida.

## Author contributions

I.K. and B.C. performed the experiments and analyzed the data. J.T. analyzed the data and developed scripts. A.M. and U.F. performed simulations of the beam shaper. K.Y.H. conceived, guided and supervised the project. B.C., I.K., A.M, U.F. and K.Y.H. wrote the manuscript. The order of the first co-authors was randomly decided.

